# Structures of metabotropic GABA_B_ receptor

**DOI:** 10.1101/2020.04.15.004267

**Authors:** Makaía M. Papasergi-Scott, Michael J. Robertson, Alpay B. Seven, Ouliana Panova, Jesper M. Mathiesen, Georgios Skiniotis

**Affiliations:** Department of Molecular and Cellular Physiology, Stanford University School of Medicine, Stanford, CA 94305, USA; Department of Structural Biology, Stanford University School of Medicine, Stanford, CA 94305, USA; Department of Drug Design and Pharmacology, Faculty of Health and Medical Sciences, University of Copenhagen, Copenhagen, Denmark

## Abstract

GABA (γ-aminobutyric acid) stimulation of the metabotropic GABA_B_ receptor results in prolonged inhibition of neurotransmission that is central to brain physiology. GABA_B_ belongs to the Family C of G protein-coupled receptors (GPCRs), which operate as dimers to relay synaptic neurotransmitter signals into a cellular response through the binding and activation of heterotrimeric G proteins. GABA_B_, however, is unique in its function as an obligate heterodimer in which agonist binding and G protein activation take place on distinct subunits. Here we show structures of heterodimeric and homodimeric full-length GABA_B_ receptors. Complemented by cellular signaling assays and atomistic simulations, the structures reveal an essential role for the GABA_B_ extracellular loop 2 (ECL2) in relaying structural transitions by ordering the linker connecting the extracellular ligand-binding domain to the transmembrane region. Furthermore, the ECL2 of both GABA_B_ subunits caps and interacts with the hydrophilic head of a phospholipid occupying the extracellular half of the transmembrane domain, thereby providing a potentially crucial link between ligand binding and the receptor core that engages G protein. These results provide a starting framework to decipher mechanistic modes of signal transduction mediated by GABA_B_ dimers and have important implications for rational drug design targeting these receptors.

## Introduction

Normal brain function depends on the proper balance and integration of excitatory and inhibitory neurotransmission^1^, a form of intracellular signaling mediated by the action of neurotransmitters on ion channels and GPCRs. The neurotransmitter γ-aminobutyric acid (GABA) is primarily responsible for synaptic inhibition throughout the nervous system via its actions on GABA_A_ ion channels and pre- and postsynaptic GABA_B_ receptors. Upon GABA stimulation, GABA_B_ receptors initiate intracellular signaling via the G_i/o_ class of heterotrimeric G proteins^2,3^, which act through the inhibition of adenylyl cyclase and voltage-gated Ca^2+^ channels, as well as the opening of G protein-coupled inward rectifying potassium channels^4^. Altogether, GABA_B_ signaling produces a prolonged decrease in neuronal excitability and modulates the release of neurotransmitters in the central nervous system. Abnormal execution of GABA_B_ signaling causes multiple neuropsychiatric diseases^5^, and the receptor is an attractive drug target for a range of disorders including drug addiction, pain, epilepsy, spasticity, anxiety, and gastroesophageal reflux disease^6-8^. In particular, the GABA_B_ agonist baclofen has been employed as an effective clinical therapeutic for the treatment of muscle spasticity^8,9^.

The Family C of GPCRs represents a small group of ~20 receptors that engage small molecule agonists, and includes GABA_B_, the metabotropic glutamate receptors (mGlu1-8), the calcium sensing receptor (CaSR), as well as two taste and several orphan receptors^10,11^. Family C receptors are unusual in that they operate as obligate dimers, with each subunit composed of a bi-lobed extracellular ligand-binding domain termed the Venus flytrap (VFT) and the common 7-transmembrane domain (7TM) of GPCRs connected via a linker region^12^. Crystallographic studies of VFTs from various Family C members, including GABA_B_, show that agonist binding rearranges the VFTs in a way that would bring the linker regions into close proximity^13,14^. All Family C GPCRs contain a cysteine-rich domain (CRD) within the linker region, with the exception of GABA_B_ which only has a relatively short linker. Our recent cryoEM structures of mGlu5 showed that the CRD interacts with extracellular loop 2 (ECL2) of the transmembrane region, thereby providing the rigidity required for transducing conformational changes 120Å from the ligand-binding site on the VFT to the 7TM domain where G protein activation occurs^15^. Given that GABA_B_ lacks a CRD, the structural communication between an agonist-bound VFT and the 7TM domains remains unclear.

Furthermore, unlike other Family C GPCRs, GABA_B_ is an obligate heterodimer comprised of two dissimilar subunits, GABA_B1_ and GABA_B2_, that must associate via intracellular C-terminal coil-coil domains in order to localize to the plasma membrane^16^. GABA_B_ is unique in that agonist binding occurs only on the VFT of GABA_B1_, while G protein coupling and activation occurs exclusively through GABA_B2_^17,18^. Thus, besides its pharmacological interest, GABA_B_ presents an ideal model system to study trans-activation mechanisms in the context of dimeric Family C GPCRs. These studies, however, are limited by the lack of structural information on full-length GABA_B_, restricting our ability to understand how agonist binding to GABA_B1_ results in G protein activation on the intracellular side of GABA_B2_. In the present study we employed single-particle cryo-electron microscopy (cryoEM), atomistic simulations, and cellular signaling assays to obtain structural and mechanistic insights into full-length GABA_B_ receptor complexes.

## Results and Discussion

The ligand-binding subunit GABA_B1_ has two isoforms, GABA_B1a_ and GABA_B1b_, differing only in the presence of two short consensus repeats at the N-terminus that localize the GABA_B1a_ isoform to the presynapse^19^. For structural studies, we designed recombinant constructs of human GABA_B1b_ and GABA_B2_ with a C-terminal hexa-histidine tag on GABA_B1b_ and an N-terminal Flag epitope (DYKDDDD) tag on GABA_B2_. The constructs were co-expressed in *Spodoptera frugiperda* (Sf9) insect cells, and the GABA_B_ heterodimer was purified by tandem affinity-chromatography in the presence of the inverse agonist CGP55845^20^ to aid receptor stability.

The structure of the inactive GABA_B_ heterodimer solubilized in glyco-diosgenin (GDN) detergent was determined by cryoEM at a global indicated resolution of 3.6 Å (Fig. 1, Supplemental Figs. 1 and 2a, Supplemental Table 1). The cryoEM map resolved the entire GABA_B_ apart from the C-terminal coil-coil domain that appears to have a flexible disposition in relation to the transmembrane regions. Additionally, we obtained a focused map of the extracellular region at a resolution of 3.5 Å, which provided improved density for the linker regions and assisted with modeling (Supplemental Fig. 1). The asymmetric protomers, GABA_B1_ and GABA_B2_, share a similar secondary structure and arrangement but are distinguished by differential glycosylation and the presence of a well-resolved density within the ligand-binding pocket of the GABA_B1_ VFT domain that is absent in GABA_B2_ (Fig. 1, Supplemental Fig. 3, Supplemental Fig. 4a). The upper VFT lobes of GABA_B_ form a junction, while the lower VFT lobes are separated by ~20 Å (Fig. 1). To model CGP55845 into the binding pocket, we utilized GemSpot^21^, our recently developed pipeline for optimal ligand docking into electron density maps. The ligand adopts a horseshoe-like conformation, closely resembling that observed in the crystal structure of the VFT domain in complex with the inhibitor CGP54626 (PDB:4MR7)^13,20^, which differs from CGP55845 only in a substitution of an aromatic ring in place of cyclohexane. The entire ligand is confined by W65 and W278 of GABA_B1_ that form hydrophobic interactions with the chlorinated ring of CGP55845 (Supplemental Fig. 3a, b). In addition, charged phosphate and amine moieties of CGP55845 are stabilized by a pair of hydrophilic residues deep within the VFT clamshell. Curiously, we observe a small spherical density in GABA_B1_ that appears to be interacting with the backbone carbonyl of W65 in addition to several anionic groups and a tyrosine, raising the possibility of a bound divalent cation in that location (Supplemental Fig. 3c). The presence of a metal ion would be consistent with the observation that calcium and other divalent ions affect ligand affinity^22,23^, and examination of the deposited scattering factors for the high-resolution VFT crystal structure (PDB:4MR7) reveals positive difference density also consistent with a cation at this site^13^.

**Figure 1.**
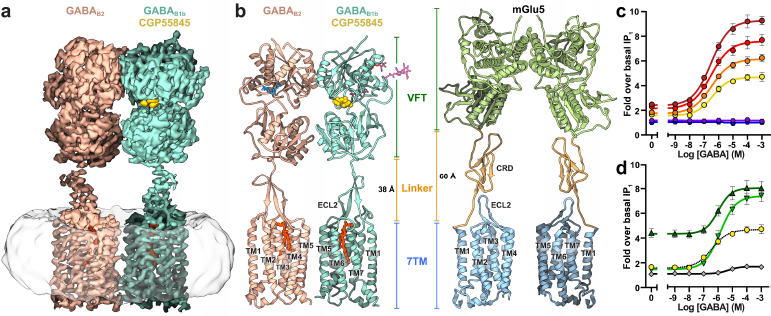
CryoEM map and model of the full-length GABA_B_ receptor heterodimer in inactive state. **a**, CryoEM map of the GABAB receptor, consisting of GABAB1b (teal) and GABAB2 (tan), within a GDN micelle (white) and harboring bound CGP55845 (yellow) and phospholipids (orange). **b**, Ribbon diagram comparing the structure of GABAB (left) with that of mGlu5 (PDB:6N52)^15^ (right). GABAB is colored analogously to corresponding EM densities in panel **a**, except for modeled N-linked glycosylation (blue and pink). The mGlu5 structure is colored by region: VFT (green), linker (orange), 7TM and ECL2 (blue). **c**, GABA efficacy increases with higher levels of co-transfected GABAB1 and GABAB2 plasmid DNA (0.8 ng, yellow; 1.6 ng, orange; 3.1 ng, red; 6.3 ng, dark red; 6.3ng GABAB1 only, purple; 3.6 ng GABAB2 only, blue). **d**, Shortening of the ECL2 of either GABAB1 (dark green) or GABAB2 (light green) co-expressed with each wild-type partner protomer increases efficacy of GABA stimulated response when compared to co-transfected wild-type receptors (yellow) of similar surface expression (Supplemental Fig. 6a), but ECL2 shortening of both protomers (grey) inhibits activity. Data in **d** represent mean ± S.E.M. from at least five independent experiments.

**Figure 2.**
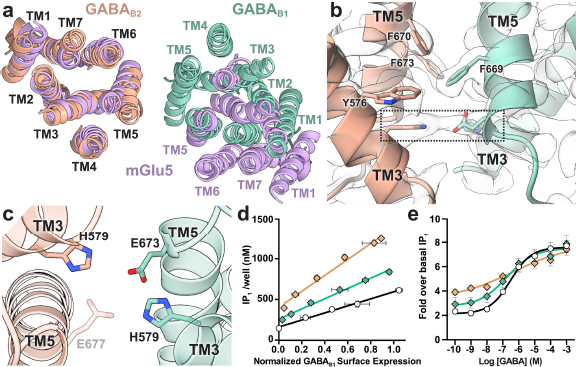
TM3 and TM5 stabilize an inactive-state dimer interface of GABA_B_ 7TM domains. **a**, Inactive-state GABA_B_ 7TMs (GABA_B1_: teal, GABA_B2_: tan) are in closer proximity compared to the 7TMs of inactive mGlu5 (purple) as shown in a top-down view. **b**, The interface between GABA_B1_ and GABA_B2_ forms a stable connection stabilized through hydrophobic interactions along TM5 helices and by polar residues on the intracellular side of TM3 and TM5 (boxed region). **c**, View of the polar interaction residues from the intracellular side of the receptor. GABA_B2_ E677 is shown for perspective; we note, however, that map density corresponding to GABA_B2_ E677 was insufficient for high-confidence modeling of its side chain. IP_1_ accumulation assays illustrate differences in GABA_B_ TM3/5 mutants’ basal activity, **d**, and GABA response, **e**. Data in **d** are representative of one experiment performed in triplicate and repeated independently at least three times with similar results. Data in **e** represent mean ± S.E.M. from at least three independent experiments.

**Figure 3.**
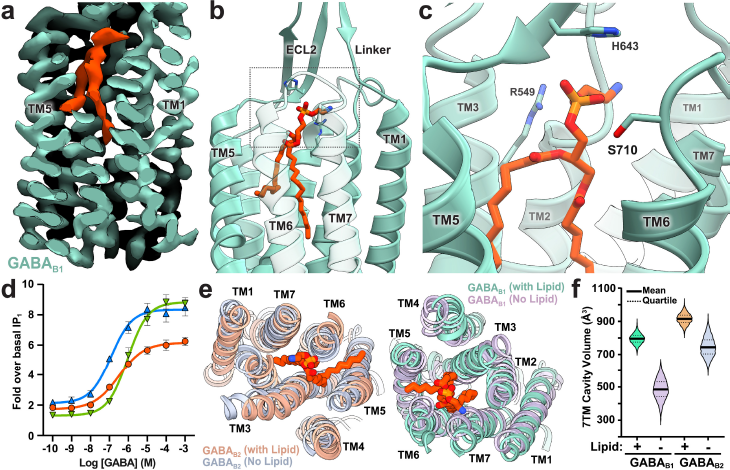
Phospholipid binds within the transmembrane cores of GABA_B_. **a**, Side view of EM map clipped to show the location of phospholipid within GABAB1 of the heterodimer. Ribbon representation of PE within GABA_B1_, **b**, with boxed region presented in panel **c** to show the structural environment of the headgroup. **d**, GABA concentration response for wild-type GABA_B_ (red), and mutants designed to displace phospholipid (GABA_B1_ L553W and GABA_B2_ L560W, green), or de-stabilize the phospholipid headgroup (GABA_B1_ R549A and GABA_B2_ R556A, blue). **e, f**, Molecular dynamics simulations show a collapse of the transmembrane cavity when lipid is absent. **e**, Representative top-down view of the GABA_B_ ribbon and stick model from molecular dynamics simulations at 200 ns. **f**, Violin plot of cavity volume ensemble data distribution over a 200 ns time course, for 3 simulations and 22,500 data points per condition. Data in **d** represent mean ± S.E.M. from at least four independent experiments.

**Figure 4.**
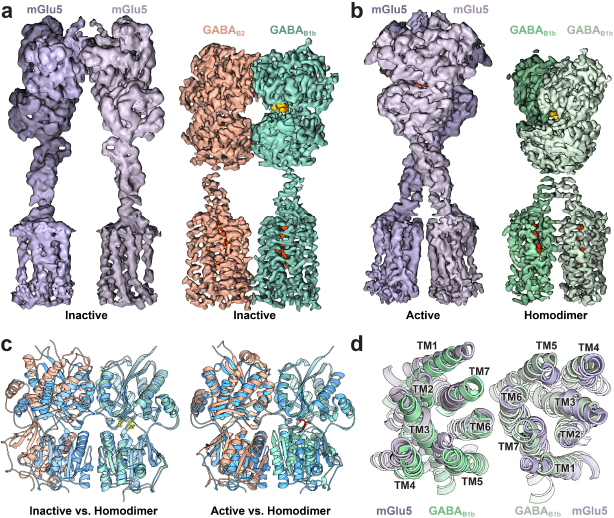
Inhibitor-bound GABA_B1_ homodimers adopt similar overall architecture and transmembrane interface as active mGlu5. **a**, EM maps of inactive mGlu5 (purple, EMD-0346) and GABA_B_ heterodimer (tan and teal); and **b**, active mGlu5 (purple, EMD-0345), and homodimeric GABA_B1_ (green). **c**, VFT overlay of crystal structures of GABA_B_ heterodimer (tan, teal) in inactive-state (left) and active-state (PDB:4MS4)^13^ (right) with the model of GABA_B1_ homodimer (blue). **d**, Top-down view of superposed 7TM domains of agonist-bound mGlu5 (purple) and GABA_B1_ homodimer (green).

Inactive GABA_B_ assumes a similar overall morphology to the apo-state structure of mGlu5^15^, while the most substantial differences arise within the linker region (Fig. 1b). Bridging the VFT and 7TM domains, an ~20 residue linker forms a β-sheet in conjunction with the extracellular loop 2 (ECL2). Notably, the length of the GABA_B_ receptor ECL2 is nearly twice that of mGlu5 (Fig.1, Supplemental Fig. 4b). The GABA_B_ cryoEM structure reveals that in the absence of CRDs, the β-sheet structure of the GABA_B_ linker in complex with ECL2 confers the structural rigidity necessary for signal transduction. Molecular dynamics simulations support the rigidity of the β-sheet formation, as after 200 ns in all simulations the linker and ECL2 continued to adopt a stable structure even in the absence of VFT domains (Supplemental Fig. 5). To further examine the involvement of the ECL2 β-sheet structure in receptor activation, we employed a functional assay with a chimeric Gα_o/q_, whereby the G_i/o_-coupled GABA_B_ receptor can couple to the PLC pathway and induce IP_3_ formation^3,24^. Accumulation of the downstream metabolite IP_1_ by LiCl was measured by an established assay^25^, thereby monitoring GABA-stimulated and basal receptor activity (Fig. 1c). The expression of GABA_B_ subunits with ECL2 deletion was generally inhibitory to the proper trafficking of the receptor to the cell surface and we thus normalized the amount of transfected DNA to obtain similar expression levels between constructs (Supplemental Fig. 6a). The deletion of the GABA_B1_ ECL2 loop (Δ627-634) produced an increase in basal activity but did not affect GABA efficacy when expressed with wild-type GABA_B2_ (Fig. 1d). These findings indicate that abrogating the ECL2-linker allows flexibility in the GABA_B1_ VFT relative to the rest of the receptor dimer, resulting in activation through the GABA_B2_ VFT/7TM route in the absence of agonist^26^. In contrast, the deletion of only the GABA_B2_ ECL2 (Δ631-638) did not affect the basal receptor activity but produced an increase in GABA efficacy and a decrease in GABA potency when compared to wild-type receptor expressed to similar levels at the cell surface, indicating that the structural rigidity of the GABA_B2_ linker region may have a partial inhibitory role (Fig. 1d, Supplemental Fig. 6a). When both receptors contain a truncated ECL2, we observed a decrease in GABA efficacy, suggesting that at least one VFT domain must be structurally coupled through the extended ECL2/linker for full activation (Fig. 1d). Collectively, these data support a bimodal transactivation mechanism of GABA_B_ in which agonist binding on the GABA_B1_ receptor can proceed from the GABA_B1_ VFT down to the GABA_B1_ 7TM region to enact changes in GABA_B2_ that promote G protein activation, and also activate the GABA_B2_ 7TM directly through the GABA_B2_ VFT domain.

Besides the VFTs and the C-terminal coiled-coil, we observe that inactive GABA_B_ forms an additional dimer interface involving interactions between TM3 and TM5 from each monomer of the heterodimer (Fig. 1, Fig. 2). This arrangement is in contrast to the structure of inactive mGlu5 receptor where the 7TM domains are separated by 16Å at their nearest point (Figs. 1b, 2a)^15^. The inactive GABA_B_ 7TM interface is formed by ionic interactions between residues on the intracellular side of TM3 and TM5 of each receptor with further stabilization through aromatic residues along the same helices (Fig. 2 b, c). Specifically, H572 (TM3) and E673 (TM5) on GABA_B1_ are in interacting proximity to H579 (TM3) and E677 (TM5) of GABA_B2_. To probe the significance of these interactions, we mutated the TM3/TM5 interface, and performed IP_1_ accumulation assays as outlined above (Fig. 2d, e, Supplemental Fig. 6). Notably, mutation of either H579 or E677 on GABA_B2_ or mutation of both E673 and H572 on GABA_B1_ increased the basal activity of the receptor, suggesting that the TM3/TM5 interface is inhibitory to signaling in the absence of agonist, which further supports the transactivation mechanism described above (Fig.2, Supplemental Figs. 6b-d). When expressed at the cell surface alone, the H579/E677 double mutant of GABA_B2_ exhibited a slight increase in basal activity compared to the silent GABA_B2_ (Supplemental Fig. 6d). These findings further support a role of an intra-protomer H579/E677 interaction in GABA_B2_ to assist stabilizing an auto-inhibited state of the GABA_B2_ protomer, and are in line with the lack of constitutive activity by GABA_B2_ ECL2 (Δ631-638).

The most unexpected observation in the transmembrane region of both monomers of GABA_B_ is a wishbone-shaped density occupying the extracellular half of each 7TM core. The bifurcated density, which is better resolved within the GABA_B1_ helical bundle, corresponds to a phospholipid (Fig. 3, Supplemental Fig. 2). Considerations of the size and shape of the density, surrounding amino acid environment within the core of the GABA_B_ 7TM, and known composition of phospholipids in Sf9 insect cells, enable us to deduce with relative confidence that the bound phospholipid in GABA_B1_ and likely GABA_B2_ corresponds to phosphatidylethanolamine (PE) (Fig. 3, Supplemental Fig. 7). The observation of PE within the transmembrane core is particularly intriguing, as no other GPCR has been shown to incorporate a two-chained phospholipid within this space. Although a lipid-activated subfamily within class A GPCRs exists, known ligands are limited to single acyl-chain lipids, eicosanoids, and sterols^27^. Those receptors are characterized by the formation of tightly structured extracellular loops preventing ligand access from the extracellular space^27^, as also observed in the cryoEM structure of GABA_B_. Notably, residues in the extracellular loops, including ECL2, and TM3 of GABA_B_ coordinate the polar head group of PE resulting in a ‘lid’ over the lipid. The remaining hydrophilic atoms in the lipid appear solvent-exposed, whereas the lipid tails are buried deep into the hydrophobic portion of the transmembrane cavity (Fig. 3b, c). Additionally, a conserved TM6 tryptophan, which acts as an activating “toggle-switch” in other Family C GPCRs^28,29^, is replaced by cysteine residues in both GABA_B1_ and GABA_B2_ receptors (Supplemental Figs. 4c, d). This replacement is essential, as any probable tryptophan rotamer at that position would clash sterically with the bound phospholipid.

The presence of lipid in the 7TM core suggests it may have a physiological role in the structural and functional integrity of the transmembrane bundles, occupying a region that corresponds to the ligand binding site in other GPCR classes (Supplemental Fig. 4). Moreover, given the crucial role of ECL2 in coordinating the linker connecting the VFT domains to the 7TM regions, it is conceivable that interactions of the phospholipid with the ECL2 have both an essential role in increasing the stability of the linker region and also participate in relaying structural transitions from the VFTs to the 7TM core. Notably, mutations designed to displace lipid tails in the 7TM core, GABA_B1_ L553W and GABA_B2_ L560W, decrease basal activity and potency of GABA while increasing GABA efficacy, whereas mutation of the arginine coordinating the head group of the phospholipids, GABA_B1_ R549A and GABA_B2_ R556A, results in increases in both GABA_B_ basal activity and receptor response to GABA (Fig. 3d, Supplemental Fig. 6e, f). Collectively, these results indicate that relaxation of phospholipid placement in the core affects receptor activation, and it is thus conceivable that the lipid directly modulates receptor activity as a result of ECL2 movements in native GABA_B_. To further probe the role of phospholipid we employed a range of atomistic simulations with the GABA_B_ receptor system. We initiated simulations of the GABA_B_ 7TM and linker heterodimer with the VFTs removed, both with and without lipid. The simulations without lipid typically diverged along one of two paths; the first being those simulations in which the transmembrane helices begin to collapse into the core with mean decreases in cavity volume of ~30% (Fig. 3, Supplemental Fig. 5), suggesting a strongly coupled interplay between the phospholipid and 7TM core. Alternatively, multiple simulations that started without lipid in the TMs showed lipid tails from the bilayer entering the receptor hydrophobic cavity below residue Y661, at a site akin to where the tail protrudes in our cryoEM structure. Remarkably, peripheral lipid tail insertion in simulations correlated with circumvention of the 7TM cavity collapse (Supplemental Fig. 5). In one simulation, after 200 ns we observed a lipid that had inserted both of its hydrocarbon tails into the receptor core. Extending the simulation by an additional 100 ns revealed the headgroup has moved over the top of the receptor, indicating lipid entry may occur as a stepwise process, with lipid tail entry sliding between TM5 and TM6 and followed by the entire lipid (Supplemental Fig. 5). Although it is probable that lipid insertion into the 7TM core occurs concurrently with helix insertion into the membrane during protein folding, these results suggest a clear tendency for phospholipids to insert at that position in a mature receptor.

Pharmaceuticals targeting the GABA_B_ receptor and other Family C GPCRs are proposed to function either at the orthosteric ligand binding site or allosterically within the transmembrane core, similar to class A orthosteric ligands. However, the presence of a phospholipid within the transmembrane core of GABA_B_ subunits would appear to occlude the binding of allosteric modulators analogous to those used to target other Family C receptors^30^. Given this observation, future developments in targeting the GABA_B_ receptor will benefit from a clearer understanding of the environment of the transmembrane core, as a potential allosteric modulator would need to either displace the core lipid or bind at an alternative site.

While a pure GABA_B_ heterodimer was obtained by tandem affinity purification of receptor constructs with distinct tags, when using non-distinct tags to purify co-expressed GABA_B1_ and GABA_B2_ constructs we found that, in addition to the GABAB1/B2 heterodimer, we also purified a significant fraction (> 40%) of GABA_B1_ homodimers. In cells, an ER retention signal within the GABA_B1_ coil-coil domain prevents homodimers from reaching the plasma membrane. Thus, the presence of GABA_B1_ homodimer in our preparation is likely due to the persistence of internal membranes during the purification procedure. On the other hand, multiple studies have suggested a physiologic role for GABA_B1_ homodimers within some cell types of the nervous system and gastrointestinal tract that express GABA_B1_ isoforms in the absence of GABA_B2_^31-36^. Although it is yet unclear how homodimeric GABA_B1_ complex functions in the absence of GABA_B2_ G protein coupling, we sought to obtain further mechanistic insights into the GABA_B_ system and obtained the structure of the GABA_B1b_ homodimer at a global indicated resolution of 3.2 Å (Fig. 4, Supplemental Figs. 2 and 8, Supplemental Table 1).

The structure of the GABA_B1_ homodimer shows that both VFTs are liganded and the two receptor monomers assume a roughly 2-fold symmetric arrangement. Notably, the VFT domains adopt the same overall conformation as the crystal structure of agonist-bound GABA_B_ VFTs (PDB:4MS4) (Fig. 4c)^13^. Furthermore, the relative arrangement of the VFT and transmembrane regions are strikingly similar to the activated state of near full-length mGlu5^15^. Thus, the conformation observed in the GABA_B1_ homodimer may provide valuable hints into the architecture of the active state GABA_B_ receptor (Fig. 4). Compared to the heterodimer, the 7TM domains in the GABA_B1_ homodimer are rotated to form an interface between the TM6 helices of each monomer. This observation is in agreement with cross-linking experiments that identified TM6-TM6 as the 7TM interaction interface upon agonist-induced stimulation of the heterodimer^37^, and agrees with structural rearrangements of mGlu5 upon agonist stimulation^15^. Along the same lines, GABA_B_ chimera studies show that replacement of the GABA_B2_ VFT with that of GABA_B1_ results in increased constitutive activity of the receptor that is not additionally stimulated by GABA^38^. Thus, the relative positioning of the VFT domains, as observed in the structure of the GABA_B1_ homodimer, appears to be sufficient to allow stabilization of an active conformation, presumably through the rigid linkage of VFT to 7TM domains to reorient the helical interface.

Collectively, our cryoEM structures, cellular signaling assays, and atomistic simulations provide crucial insights into mechanistic aspects of GABA_B_ signaling. The extended ECL2 and its interaction with the linker region appears to compensate for the lack of CRD in GABA_B_ compared to all other Family C receptors, and thus also rigidly transduce conformational changes from the VFT to the 7TM domains. The presence of phospholipid occupying the extracellular half of the 7TM core, which has not been previously observed for any GPCR, appears important for the structural integrity of the transmembrane domains in both GABA_B_ subunits, while also structurally coordinating the critical ECL2 region. Our findings support a model in which agonist binding to GABA_B1_ results in VFT dimer compaction that reorients the rigidly structured linker/ECL2. Such conformational change would drive the 7TM domains to twist away from an auto-inhibited state mediated by the inactive TM3/TM5 interface, thereby forming a new interface mediated by the TM6 helices. Although currently unclear, the activating transitions within the GABA_B2_ 7TM are likely mediated both through the newly formed TM6/TM6 interface and propagation through its own ECL2, as supported by our functional assays. An intriguing possibility is that the phospholipid may act as a sensor of changes in VFT and ECL2 conformations that would result in activating transitions within the 7TM domain of GABA_B2_ to prime it for G protein engagement. Addressing these questions will require further structural studies of active state heterodimer alone and in complex with G protein. The present work, along with our recent studies on mGlu5, form a starting structural framework to decipher the enigmatic signal transduction mechanism of GABA_B_ and Family C GPCRs in the context of full-length receptors.

## Materials & Methods

### Cloning

The cDNA clone for human GABA_B2_ receptor (Accession: NM_005458) in pcDNA3.1^+^ was obtained from the cDNA Resource Center (www.cdna.org); and the cDNA clone for human GABA_B1b_ was purchased from Horizon Discovery (Accession: BC050532, Clone ID: 5732186). Primers were designed to include a hemagglutinin (HA) signal sequence^39^ in the place of authentic signal sequences of each receptor, thus removing the first 29 residues of GABA_B1_ and 41 residues of GABA_B2_. The following primers (Integrated DNA Technologies) were used to sub-clone GABA_B_ constructs into the pFastBacDual vector (Invitrogen) with N-terminal Flag epitope (DYKDDDD) following the HA tag and/or C-terminal hexahistidine (His6) tags: EcoRI-HA-Flag-GABA_B1,_5’-GCGCGCGAATTCATGAAGACGATCATCGCCCTGAGCTACATCTTCTGCCTGGTGTTCGCCGATTACAAGGACGACGATGACAAGTCCCACTCCCCCCATCTCCCG-3’; GABA_B1_-His6-SalI, 5’-GCGCGCGTCGACTTAATGATGATGATGATGGTGCTTATAAAGCAAATGCAC-3’; GABA_B1_-SalI 5’-GCGCGCGTCGACTTACTTATAAAGCAAATGCAC-3’; EcoRI-HA-FLAG-GABA_B2_, 5’-GCGCGCGAATTCATGAAGACGATCATCGCCCTGAGCTACATCTTCTGCCTGGTGTTCGCCGATTACAAGGACGACGATGACAAGTGGGCGCGGGGCGCCCCC-3’; GABA_B2_-His6-SalI, 5’-GCGCGCGTCGACTTAATGATGATGATGATGGTGCAGGCCCGAGACCATGAC-3’; GABA_B2_-SalI, 5’-GCGCGCGTCGACTTACAGGCCCGAGACCATGAC-3’. To generate GABA_B_ mutants PCR reactions were conducted with either PfuTurbo or Q5 polymerase using the following pairs of primers and pcDNA3.1+ containing either HA-FLAG-GABA_B1b_ or HA-HA-GABA_B2_: GABA_B1b_ H572A, 5’-GGTGGGTCGCCACGGTCTTC-3’ and 5’-GAAGACCGTGGCGACCCACC-3’; GABA_B1b_ E673A, 5’-CTTGCTTATGCTACCAAGAG-3’ and 5’-CTCTTGGTAGCATAAGCAAG-3’; GABA_B2_ H579A, 5’-CTGGAGAGTCGCTGCCATCTTCAA-3’ and 5’-TTGAAGATGGCAGCGACTCTCCAG-3’; GABA_B2_ E677A, 5’-CTTAGCTTGGGCTACCCGCAAC-3’ and 5’-GTTGCGGGTAGCCCAAGCTAAG-3’; GABA_B1_ (Δ627-634), 5’-CTTGGCAAATGTCTCAATGGTC-3’ and 5’-GTCTCTATTCTGCCCCAGC-3’; GABA_B2_ (Δ631-638), 5’-ATCTCCATCCGCCCTCTCC-3’ and 5’-CATGCTGTACTTCTCCACTG-3’; GABA_B1_ R549A, 5’-CTGCCAGGCCGCCCTCTGGCTCCTG-3’ and 5’-ACGAAAGGGAACTGG-3’; GABA_B2_ R556A, 5’-GCACCGTCGCTACCTGGATTCTC-3’ and 5’-AAA GTG TTT CAA AGG-3’; GABA_B1_ L553W, 5’-CTCTGGCTCTGGGGCCTGGGCTTTAG-3’ and 5’-GCGGGCCTGGCAGACG-3’; GABA_B2_ L560W, 5’-GGACCTGGATTTGGACCGTGGGCTAC-3’ and 5’-TGACGGTGCAAAGTG-3’.

### Expression and Purification

*Spodoptera frugiperda* (Sf9) insect cells (Expression Systems) were co-infected at a density of ~2.0 × 10^6^ cells/mL with HA-FLAG-GABA_B2_ baculovirus and either HA-FLAG-GABA_B1b_-His6 or HA-GABA_B1b-_His6 baculovirus at a multiplicity of infection (M.O.I.) between 3.0 – 5.0. During expression, cells were treated with 5 μM CGP55845 (Hello Bio, Inc.). At 48 hours post-infection cells were harvested by centrifugation, washed once with phosphate-buffered saline containing protease inhibitors (leupeptin, soybean trypsin inhibitor, N-*p*-Tosyl-L-phenylalanine chloromethyl ketone, Tosyl-L-lysyl-chloromethane hydrochloride, phenylmethylsulfonyl fluoride, aprotinin, bestatin, pepstatin) and 5 μM CGP55845. Cell lysis was achieved through nitrogen cavitation in buffer containing 20 mM HEPES, pH 7.5, 150 mM NaCl, 1 mM EDTA, 10 μM CGP55845, 2 mM MgCl_2_, nuclease, and protease inhibitors. The whole-cell lysate was centrifuged at 1,000 xg to remove nuclei and unbroken cells. The supernatant was centrifuged at 100,000 xg to isolate the membrane fraction. Membranes were resuspended by Dounce homogenization in buffer containing 20 mM HEPES, pH 7.5, 150 mM NaCl, 2 mM MgCl_2_, 1 mM EDTA, 2 mg/mL iodoacetamide, 10 μM CGP55845, 1% n-Dodecyl β-D-maltoside (DDM), 0.2% Sodium cholate, 0.2% Cholesterol hemisuccinate (CHS), nuclease, and protease inhibitors. Solubilized membranes were clarified by centrifugation at 100,000 xg, and the supernatant was loaded onto a pre-equilibrated column of anti-DYKDDDDK G1 affinity resin (Genscript). The resin was washed with Buffer A (20 mM HEPES, pH7.5, 150 mM NaCl, 10 μM CGP55845, and protease inhibitors) with 0.1% DDM and 0.02% CHS. Protein was eluted with Buffer A containing 0.1% DDM, 0.02% CHS, and 0.2 mg/mL DYKDDDDK peptide. The eluate was then loaded onto a pre-equilibrated Nickel-NTA column. Resin was washed with Buffer A containing 0.1% DDM, 0.02% CHS; and the buffer was exchanged in six steps to Buffer A supplemented with 0.2% GDN, 0.02% CHS, followed by a two-step exchange into Buffer A containing 0.004% GDN, 0.0004% CHS. Protein was eluted from the Ni-NTA resin with Buffer A containing 0.004% GDN, 0.0004% CHS, and 500 mM Imidazole. The resulting eluate was concentrated by centrifugal filtration with a 50 kDa molecular weight cut off, and subsequently run on a Superose 6 size exclusion column (GE Healthcare). Samples were pre-screened for sample quality by negative stain transmission electron microscopy and then immediately prepared on cryoEM grids.

### CryoEM Data Collection

For the GABA_B1b_/GABA_B2_ heterodimer, 3.5 uL of sample was applied at a concentration of 3-5 mg/mL to glow-discharged holey carbon grids (Quantifoil R1.2/1.3). The grids were blotted using an FEI Vitrobot Mark IV (Thermo Fisher Scientific) at 18 °C and 100% humidity, and plunge frozen into liquid ethane. Two data sets were used to produce the final structure. For both data collections cryoEM imaging was performed on a Titan Krios (Thermo Fisher Scientific) electron microscope equipped with a K3 Summit direct electron detector (Gatan). The microscope was operated at 300 kV accelerating voltage, at a magnification of 57,050x in counting mode resulting in a magnified pixel size of 0.8521 Å. For the first data set, movies were obtained at a dose rate of 14.19 electrons/ Å^2^/sec with defocus ranging from −1.5 to −2.7 μm. The total exposure time was 3.985 sec over 57 frames per movie stack. For the second data set, movies were obtained at a dose rate of 21.43 electrons/ Å^2^/sec with defocus ranging from −1.2 to −2.5 μm. The total exposure time was 2.996 sec including 50 frames per movie stack.

CryoEM grids for the GABA_B1b_ homodimer at a concentration 5.0 mg/mL were prepared similar to the heterodimer. CryoEM imaging was performed on a Titan Krios electron microscope equipped with a post-column energy filter and a K2 Summit direct electron detector (Gatan). The microscope was operated at 300 kV accelerating voltage, at a magnification of 47,198x in counting mode resulting in a pixel size of 1.06 Å. Movies were obtained at a dose rate of 6.212 electrons/Å^2^/sec with defocus ranging from −0.9 to −2.5 μm. The total exposure time was 8.0 sec over 40 frames per movie stack. Automatic data acquisition was performed using SerialEM^40^ for all data sets.

### Image Processing and 3D Reconstruction

Dose-fractionated image stacks were subjected to beam-induced motion correction and dose-weighting using MotionCor2^41^. Contrast transfer function parameters for each non-dose weighted micrograph were determined by Gctf^42^ for the homodimer and data set #1 of the heterodimer, and by CtfFind-4.1^43^ for data set #2 of the heterodimer. For all data sets; particle selection, 2D and 3D classification were performed on a binned dataset (pixel size 1.72Å and 4.24Å for the heterodimer and homodimer, respectively) using RELION (versions 3.0 and 3.1)^44^. The two data sets for the heterodimer were processed individually before being combined following a Bayesian polishing step. A total of 538,957 particles from 1,324 micrographs and 2,062,083 particles from 8,991 micrographs were extracted using semi-automated particle selection for the heterodimer data set #1 and #2, respectively. Both particle sets were then separately subjected to three rounds of 2D classification and two rounds of 3D classification. Particles in both sets were subjected to Bayesian polishing individually and then combined for a total of 286,140 particles. The merged dataset was fit for Ctf parameters (per particle defocus and astigmatism, per micrograph *B*-factor) and estimated for anisotropic magnification and beam-tilt. A final 3D refinement was followed by post-processing using a mask that excluded the GDN micelle density. A focused refinement was also carried out using a mask encompassing the VFT and linker regions of GABA_B_. For the GABA_B1b_ homodimer structure, a total of 2,278,113 particles were extracted from 5,602 micrographs using semi-automated particle selection. Particles were subjected to multiple rounds of 2D and 3D classification until a subset of 282,811particles were selected for the final map. The particle set underwent multiple rounds of Ctf parameter fitting and was subjected to Bayesian polishing before 3D Refinement and post-processing of the final map. UCSF Chimera^45^ was used for map/model visualization.

### Model Building

The initial model for the VFT domain was taken from the inactive state VFT crystal structure PDB:4MR7^13^ and the initial structure of the transmembrane domain of GABA_B1b_ was generated as a homology model from the inactive cryoEM structure of mGlu5 (PDB:6N52) using Schrödinger’s Prime homology modeling^15^. Both components were placed into the GABA_B_ cryoEM map using Chimera’s ‘fitin-map’ function. The linker, intracellular loops, and extracellular loops of GABA_B1_ were interactively adjusted into the EM map using Coot (version 0.8.9.1el)^46^ and the resulting model of the GABA_B1_ Linker/7TM was then used to generate a homology model of GABA_B2_ using Schrodinger’s Prime homology modeling, which was also placed into the map in Chimera. Iterative rounds of interactive model adjustment in Coot followed by real-space refinement in Phenix (version 1.17.1-3660)^47^ employing secondary structure restraints in addition to the default restraints were performed to improve the modeling. Once confidence in the sidechain placement was reached for the ligand-binding cleft on GABA_B1_, the GemSpot pipeline^21^ (Schrödinger) was used to model the inhibitor, CGP55845, into the map. After further improvement, 1-palmitoyl-2-oleoyl-sn-glycero-3-phosphoethanolamine (POPE) was modeled in the transmembrane pocket of GABA_B1_ and GABA_B2_ with the GemSpot pipeline. Final refinement was performed with Phenix^47^.

### Molecular Dynamics Simulations and Analysis

To prepare the system for molecular dynamics simulations, the low-resolution features of the map were used to manually build ICL2 into the model of the GABA_B1_/GABA_B2_ inactive heterodimer using Coot. The system was then prepared in Maestro, version 2019-4 (Schrödinger) to build any stubbed sidechains and determine protonation states. The VFTs were removed from the heterodimer to produce a truncated construct starting at residues T461 for GABA_B1b_ and T468 for GABA_B2_, thus containing only the linkers and the TM domains. The Orientations of Proteins in Membranes (OPM)^48^ webserver was used to orient the system with respect to a membrane plane and the CHARMM-GUI^49^ was employed to place the system in either a 1-palmitoyl-2-oleoyl-sn-glycero-3-phosphocholine (POPC) and cholesterol bilayer or a 3:1 POPC:1-palmitoyl-2-oleoyl-sn-glycero-3-phosphoethanolamine (POPE) and cholesterol bilayer. Approximate dimensions for the system were 105 x 105 x 110 Å for a total of 240 lipid and 7 cholesterol molecules. This bilayer was then solvated in TIP3P water with 150 mM sodium chloride ions balanced to achieve charge neutrality. POPE was used for the lipid in the TM binding sites of GABA_B1_ and GABA_B2_.

The PDB file for the full solvated system was prepared in VMD (version 1.9.3)^50^ for simulation in NAMD (version 2.13)^51^. The OPLS-AA/M^52^ force field was used for the protein, while OPLS-AA^52^ was used for the lipids, cholesterol, and ions. Disulfide bonds were placed between C546 and C644 in GABA_B1_ and C553 and C648 in GABA_B2_ and both the N- and C-termini were blocked with capping groups. NAMD was used to run molecular dynamics simulations, where all phases employed periodic boundary conditions with non-bonded interactions smoothed starting at 10 Å to 12 Å, with long range interactions treated with the particle mesh Ewald method. Systems were minimized for 2000 steps and then slowly heated in the NPT ensemble with a Langevin thermostat and a Nosé-Hoover Langevin piston barostat set at 1 atm with a period of 50 fs and a decay of 25 fs. A 2 fs time-step was used with the SHAKE^53^ and SETTLE^54^ algorithms. Heating occurred from 0 K to 310 K in increments of 20 K with 0.4 ns of simulation at each increment. Harmonic restraints of 1 kcal/mol/Å^2^ were used during heating on all non-hydrogen atoms of the protein and lipids. The system was then equilibrated with 1 kcal/mol/Å^2^ harmonic restraints on all protein and lipid non-hydrogen atoms for 10 ns followed by another 10 ns of equilibration with 1 kcal/mol/Å^2^ harmonic restraints on non-hydrogen backbone atoms. Finally, 1 kcal/mol/Å^2^ harmonic restraints were applied to only C alpha atoms for 2 ns before being stepped down to 0.5 kcal/mol/Å^2^ for 2 ns, 0.3 kcal/mol/Å^2^ for 2 ns, and then removed. The first 30 ns of unrestrained molecular dynamics were also discarded as equilibration.

All trajectories were down sampled by 10x for analysis. Cavity volume was calculated with Epock (1.0.5)^55^ in VMD^50^ on trajectories that had been aligned to either GABA_B1_ or GABA_B2_ from the starting structure. The cavity region was defined to include the binding region of the hydrophobic tails of the lipid. TM-TM distances were calculated in VMD based on the CA position of residues: 3.33, 4.50, 5.40, 6.54, and 7.28 in the Ballesteros–Weinstein^56^ numbering scheme.

### Transfection and seeding of cells for signaling assays

HEK293 cells (ATCC^®^ CRL-1573^™^) were transfected with expression vector DNAs encoding the two GABA_B_ receptor protomers and a chimeric Gα_q/o5_ subunit (five C-terminal amino acids of Gα_q_ were exchanged with those of Gα_o_) to allow the Gα_i/o_-coupled GABA_B_ receptor to activate PLC and induce IP_3_ and intracellular Ca^2+^ release^3^. Prior to transfection cells were brought into suspension by trypsinization and resuspension to 0.18 million cells/mL in growth medium (D-MEM, Gibco 10566016; supplemented with 10% Fetal Bovine Serum, Gibco 10270106; 1% Sodium Pyruvate, Gibco 11360039; 1% MEM Non Essential Amino Acids, Gibco 11140068; and 1% Penicillin-Streptomycin Solution, Gibco 15140122).

For each 1 mL of cell suspension transfected, a total of 1 µg DNA in 25 µL OptiMEM (Gibco 51985) was incubated for 20 min with a mixture of 57 µL OptiMEM and 3 µL FuGene6 (Promega E2692). After FuGene6/DNA complex formation, the mixture was added directly to the cell suspension, mixed thoroughly and cells seeded with 100 µL cell suspension in appropriate 96-well plates. Of the 1 µg DNA/mL cell suspension, the amount of expression vector DNA encoding the chimeric Gα_q/o5_ was 0.5 µg/mL cell suspension in all experiments. The amount of GABA_B_ encoding DNA was varied between 7.8 ng and 0.25 µg for each of the GABA_B_ receptor protomer DNAs depending on the assay and mutants tested. Empty vector DNA was added to give a total amount of 1 µg DNA/mL cell suspension transfected. For characterization of basal activity of the TM3/5 protomer mutants, a typical gene dose experiment was performed. DNA corresponding to 62.5 ng DNA/mL cell suspension of each of the protomers (wild-type or mutants) were mixed and serially diluted 2-fold 5 times, typically down to 3.9 ng DNA/mL cell suspension. The transfected cell suspension was seeded at 100 µg/mL both in clear poly-L-lysine coated 96-well plates for IP_1_ accumulation assays and in white poly-L-lysine coated 96-well plates for cell surface ELISA assays.

### IP_1_ accumulation assays

The IP_1_ assays for wild-type and mutant receptors was performed essentially as described^15^. Forty-eight hours after transfection, the growth medium was replaced with HBSS buffer (HBSS (Gibco 14025), 20 mM HEPES pH 7.5, 1 mM CaCl_2_, 1 mM MgCl_2_ and 0.1% BSA) supplemented with BSA to 0.5% and incubated at 37 °C for 3-4 hours.

For characterization of the TM3/TM5 protomer interface mutants for basal activity, the HBSS + 0.5% BSA buffer was replaced with 100 µL HBSS buffer, followed by addition of 50 µL HBSS buffer containing LiCl (150 mM) to give a final concentration of 50 mM LiCl. After incubation for 1 hour at 37°C the IP_1_ accumulation was stopped by addition of 40 μL CisBio IP-One Tb HTRF Kit (CisBio, 62IPAPEC) lysis buffer. The accumulated IP_1_ levels were determined according to the manufacturer’s instructions and as described^15^.

For the generation of GABA concentration response curves, the compounds were diluted in three times the final concentration in HBSS buffer containing 60 mM LiCl. The assay was started, first by replacing the HBSS + 0.5% BSA buffer with 100 µL HBSS buffer, followed by addition of 50 µL of the above compound dilutions to give a final LiCl concentration of 20 mM. The IP_1_ accumulation assay was stopped and assayed as described above after incubation for 1 hour at 37 °C. Data were calculated as the amount of IP_1_ formed per well or normalized to the basal IP_1_ level and fitted by non-linear regression using GraphPad Prism. Results and description of statistical analyses used in this manuscript are found in Supplemental Table 2.

### Cell surface ELISA assay

Surface expression levels of wild-type and mutant GABA_B_ receptors were determined using a direct enzyme-linked immunosorbent assay (ELISA) against the N-terminal GABA_B1b_ FLAG tag and the N-terminal GABA_B2_ HA-tag, as described^57^. Transfected cells were seeded in white Poly-D-Lysine-coated 96-well plates. Forty-eight hours after transfection, cells were washed once with 100 µL/well DPBS + 1 mM CaCl_2_ (wash buffer). Following fixation with 50 µL/well 4% paraformaldehyde solution for 5 minutes at room temperature, cells were washed twice with 100 µl wash buffer and blocked with 100 µL/well blocking solution (3% dry-milk, 1 mM CaCl_2_, 50 mM Tris-HCl, pH 7.5) for 30 minutes at room temperature, followed by addition of 75 µL/well HRP-conjugated anti-FLAG antibody (Sigma Aldrich, A8592), or HRP-conjugated anti-HA antibody (R&D systems HAM0601), both diluted 1:2000 in blocking solution, and allowed to incubate for 1 hour at room temperature. The plates were then washed four times with 100 µl/well blocking solution followed by four washes with wash buffer. The amount of surface expressed receptors was detected by adding 60 µL wash buffer and 20 µL HRP substrate (Bio-Rad, 170-5060) per well, incubating for 10 minutes and measuring of luminescence in an EnVision plate reader (Perkin Elmer).

## Author contributions

M.M.P-S. designed and cloned GABA_B_ constructs, expressed and purified all proteins, collected and processed cryoEM data. M.J.R. built and refined the structure from cryoEM density maps and setup, performed, and analyzed molecular simulations. A.B.S. and O.P. assisted with cryoEM data collection and processing. J.M.M. performed and analyzed cellular signaling experiments. M.M.P-S., M.J.R, J.M.M., and G.S. interpreted results. M.M.P-S., M.J.R., and G.S. wrote the manuscript with J.M.M., O.P., and A.B.S. providing input. G.S. supervised the project.

## Acknowledgements

The work is supported by National Institutes of Health (NIH) grant R01 NS092695 (G.S. and J.M.M.) and used the Extreme Science and Engineering Discovery Environment (XSEDE)^58^ resource comet-gpu through sdsc-comet allocation TG-MCB190153, which is supported by National Science Foundation grant number ACI-1548562. We thank Qianhui Qu for cryoEM advice and assistance.

---

**Supplementary Figure 1.**
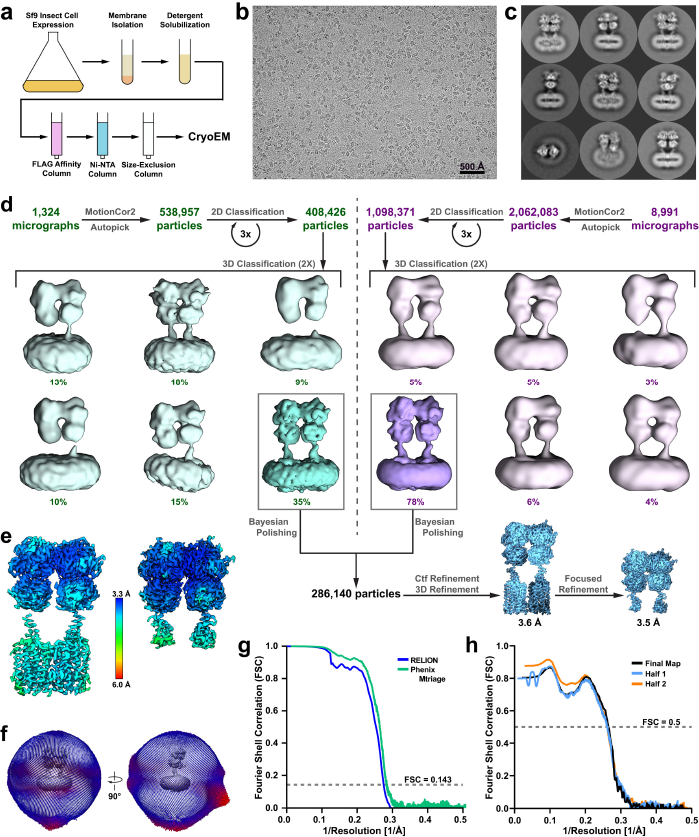
Sample preparation, cryoEM processing and reconstruction of GABA_B_ heterodimer. **a**, Purification scheme for GABA_B_. Representative cryoEM micrograph (**b**) and 2D class averages (**c**) of GABA_B_ dimers. **d**, Flow chart outlining the cryoEM processing workflow using RELION^59^, the global resolutions of the full-length structure and VFT focused structures were 3.6 Å and 3.5 Å, respectively, at 0.143 Fourier shell correlation (FSC) as calculated by RELION. **e**, Local resolution of cryoEM maps. **f**, Angular distribution of projections used in final cryoEM reconstruction. **g**, **h**, Gold standard FSC curve of half-maps calculated using RELION and Phenix Mtriage^60^ (**g**), and map-to-model validation curves generated through Phenix Mtriage (**h**).

**Supplementary Figure 2.**
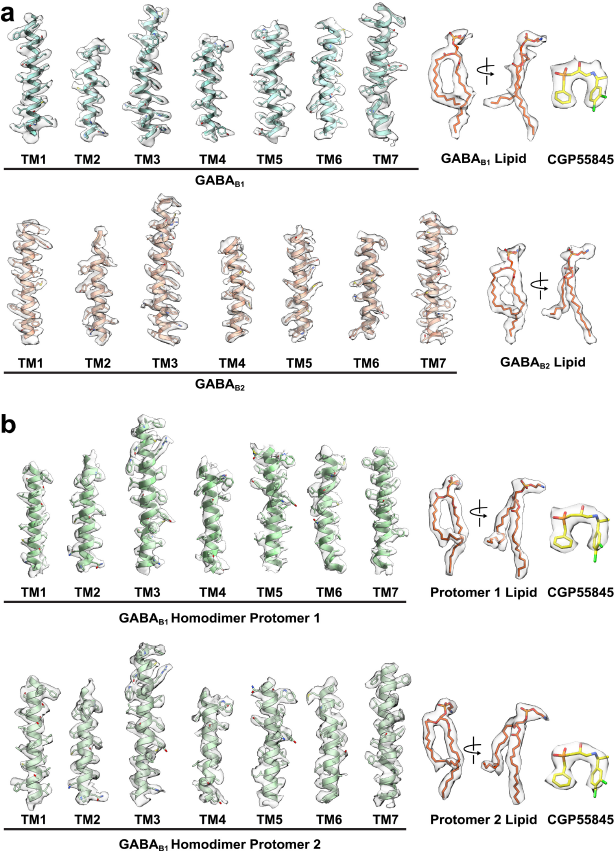
Agreement between cryoEM map and model. **a**, EM density and model for GABA_B_ heterodimer complex; transmembrane helices of GABA_B1_, transmembrane helices of GABA_B2_, bound PE, and ligand CGP55845. Densities visualized within UCSF Chimera^45^ and zoned at 2.2 with threshold set to 0.0142, with the exception of GABA_B1_-bound lipid and GABA_B1_-bound CGP55845 in which thresholds of 0.01 and 0.0189 were used, respectively. **b**, EM density and model for GABA_B1_ homodimer; transmembrane helices of both protomers, 7TM-bound PE, and ligand CGP55845. Densities were zoned at 2.2 and threshold set to 0.016, apart from CGP55845 in both protomers in which a threshold of 0.03 was used.

**Supplementary Figure 3.**
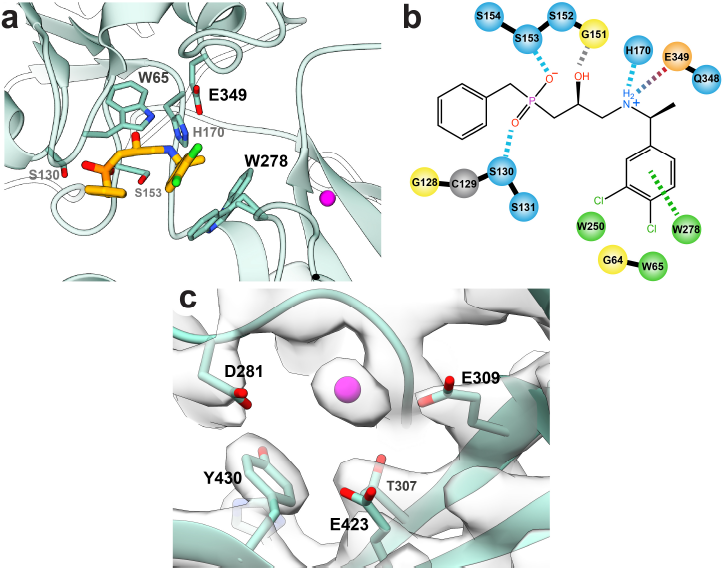
Binding of CGP55845 and cation to GABA_B1_ VFT domain. **a**, Model of CGP55845 within the VFT of GABA_B1b_. **b**, Schematic of interacting residues on GABA_B1b_ with the inhibitor, CGP55845. GABA_B1b_ residues S153 and S130 form hydrogen bonds with oxygen atoms of the phosphate group, while H170 and E349 form a hydrogen bond and a salt bridge with the amine group of the ligand, respectively. π-π stacking occurs between the chlorinated ring structure of CGP55845 and W278, while W65 provides hydrophobic packing on the opposing side of the ring. S130, H170, E349, and W65 are all substantially different residues in GABA_B2_, precluding ligand binding. Residues are color-coded corresponding to their properties: light blue, hydrophilic; orange, anionic; green, hydrophobic; yellow, glycine; and grey, cysteine. Interaction lines are also color-coded according to their type: light blue, side-chain hydrogen bonding; grey, backbone hydrogen bonding; blue-red gradient, salt-bridge; and green, π-π stacking. **c**, Spherical density surrounded by anionic residues within the VFT supports a cation (magenta) at that site.

**Supplementary Figure 4.**
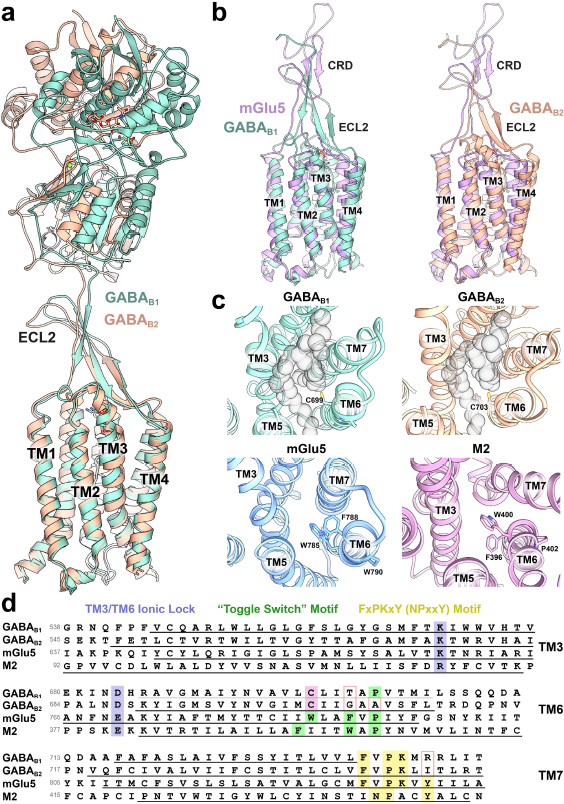
Comparison of structures across GPCR classes. **a**, GABA_B_ protomers share similar secondary and overall structure. **b**, Comparison of mGlu5 and GABA_B_ 7TM and ECL2/linker shown from side view. **c**, Top-down view of GABA_B1_, GABA_B2_, mGlu5 (PDB: 6N52), and class A M2 acetylcholine receptor (PDB:3UON) with side-chains corresponding to the “toggle-switch” motif shown. Phospholipid space-filling model is included in gray within GABA_B1_ and GABA_B2_. **d**, Sequence alignment of human GABA_B_ receptors with mGlu5 and M2 receptors comparing canonical GPCR activation motifs: TM3-TM6 ionic lock (blue), “toggle switch motif” (green), FxPKxY motif (yellow). Sequences are aligned to motifs within each TM helix and transmembrane helical secondary structure is underlined. Residues in GABA_B_ sequences differing from canonical motifs are outlined in pink. The cysteine residue that replaces the “toggle-switch” tryptophan is highlighted in pink.

**Supplementary Figure 5.**
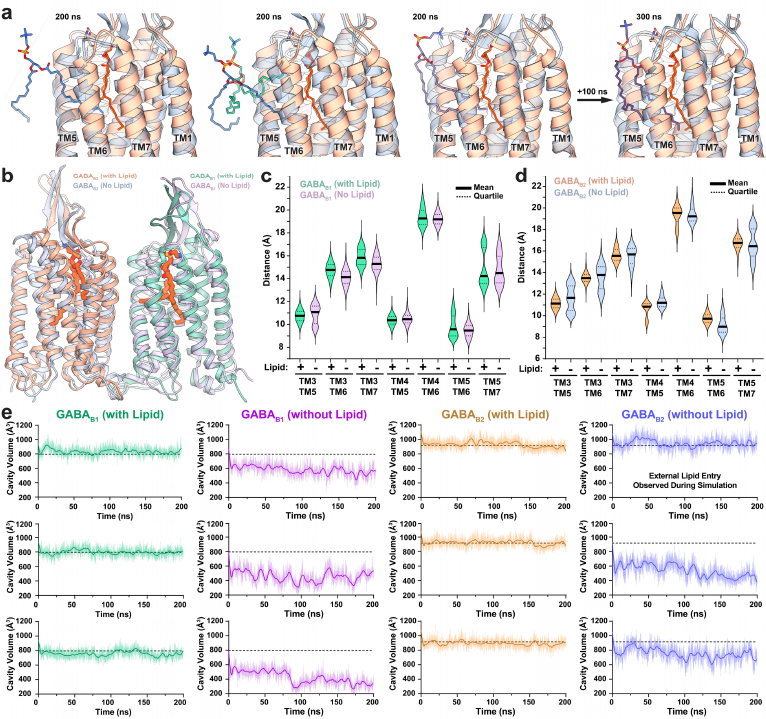
Atomistic simulations of phospholipid structural stabilization and entry. **a**, The results of three out of seven total simulations of GABA_B_ 7TM and linker in the absence of core-bound lipid after 200 ns. The results show the extent of lipid (green, purple) entry in GABA_B2_ (grey) in the simulations versus the experimental structure (tan) with PE (orange). One of the trajectories was extended an additional 100 ns (rightmost panel) and lipid entry was observed to progress towards the core. **b**, Representative side view of the GABA_B_ ribbon and stick model from simulations at 200 ns showing rigidity of ECL2 β-sheet structure even in absence of VFT domains. **c**, **d**, Violin plots of ensemble distances between GABA_B1_ (**d**) and GABA_B2_ (**e**) TM helices in simulations with and without core-bound lipid. Distances were measured from the Cα atoms of the following residues in GABA_B1_: L550 (TM3), W611 (TM4), L653 (TM5), M707 (TM6), A716 (TM7); and the following residues in GABA_B2_: T557 (TM3), W615 (TM4), L657 (TM5), F711 (TM6), Q720 (TM7). Simulations were run in triplicate over 200ns and violin plots represent 25,000 data points per condition. **e**, 7TM cavity volume measured over time for individual simulations. Average cavity volume of phospholipid-bound receptor is shown as dashed line, thick lines indicate rolling averages of 5.33 ns, and thin lines represent raw data.

**Supplementary Figure 6.**
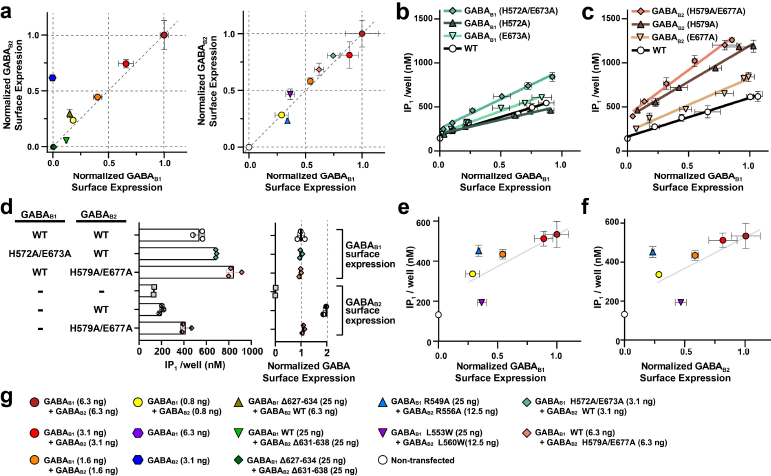
Functional analysis of GABA_B_ mutants. **a**, Comparative normalized surface expression levels of constructs assessed by FLAG-tagged GABA_B1_ and HA-tagged GABA_B2_ surface expression. **b**, **c**, Mutations of ionic residues forming the interface of GABA_B1_ (**b**) and GABA_B2_ (**c**) result in increased constitutive activity of the receptor. **d**, GABA_B2_ H579A/E677A expressed without GABA_B1_ shows a moderate increase in basal activity over wild-type GABA_B2_ expressed alone. **e, f**, Mutation of lipid coordinating residues (blue triangle) increase the constitutive activity of GABA_B1_ (**e**) and GABA_B2_ (**f**), while mutations displacing the lipid tails from the 7TM core (purple triangle) result in decreased basal activity of the receptor when compared to wild-type receptor of similar receptor surface expression. **g**, Key to colors and symbols used in panels **a**, **e**, and **f** with DNA transfection amounts indicated in parenthesis. Data are representative of one experiment performed in triplicate and repeated independently at least three times with similar results.

**Supplementary Figure 7.**
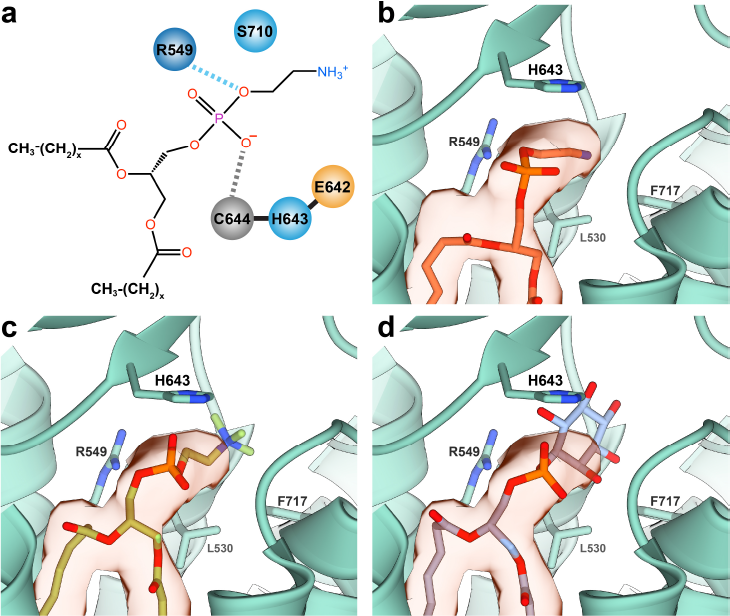
Modeling of phospholipid into GABA_B_. **a**, Schematic of GABA_B1_ residues interacting with the polar head group of phosphatidylethanolamine (PE). The terminal amine (-NH^3+^) forms a salt bridge with residue D714^ECL3^, while the phosphate group is coordinated by S710^ECL3^, R549^TM3^, and the backbone nitrogen of C644^ECL2^. **b-d**, As our receptor purification did not contribute additional lipid, we considered the known lipid composition of Sf9 insect cells. Four primary phospholipids are present in Sf9 insect cells: phosphatidylcholine (43%), phosphatidylethanolamine (32%), phosphatidylinositol (23%), and cardiolipin (4%)^61^. It was apparent from the map that the lipid had only two carbon chains, immediately excluding cardiolipin as it has four hydrocarbon tails. A comparison of the map and the binding site residues led to the decision to model PE (**b**) into the pocket using the GemSpot pipeline^21^, which produced good cross-correlation with favorable interactions. To further confirm our selection, analysis of overlays of phosphatidylcholine (**c**) and phosphatidylinositol (**d**) over the docked model revealed phosphatidylcholine is unlikely given that the interactions with the cation appear to be primarily salt bridges, rather than the cation-π interactions that more commonly coordinate choline in proteins^62^. Although phosphatidylinositol may make favorable interactions, our map does not appear to support such a large moiety in the head group position. Thus, PE is the most likely lipid to reside in the structure and was thus used in the models.

**Supplementary Figure 8.**
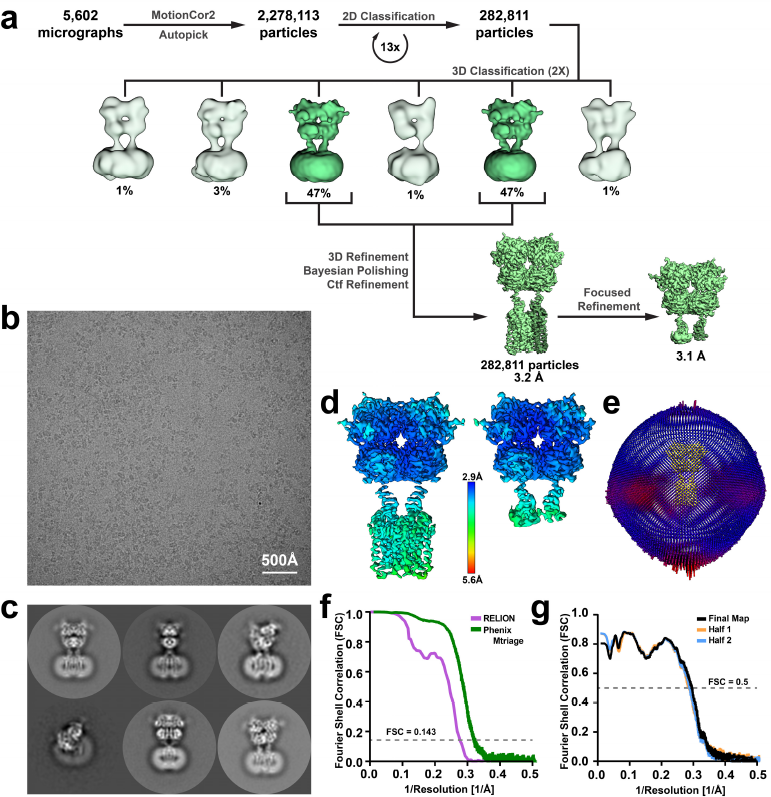
CryoEM processing workflow of GABA_B1b_ homodimer. **a**, Flow chart outlining the cryoEM processing of the GABA_B_ homodimer. **b**, **c**, Representative micrograph (**b**) and 2D class averages (**c**). **d**, Local resolution of cryoEM maps. **e**, Angular distribution of projections employed in the final cryoEM reconstruction. **f**, **g**, Gold standard FSC curve of half-maps calculated using RELION and Phenix Mtriage^60^ (**f**), and map-to-model validation curves generated through Phenix Mtriage (**g**). Global indicated resolutions of the full-length structure and VFT structure were 3.2 Å and 3.1 Å, respectively, at 0.143 Fourier shell correlation (FSC) as calculated by RELION.

**Supplemental Table 1.**
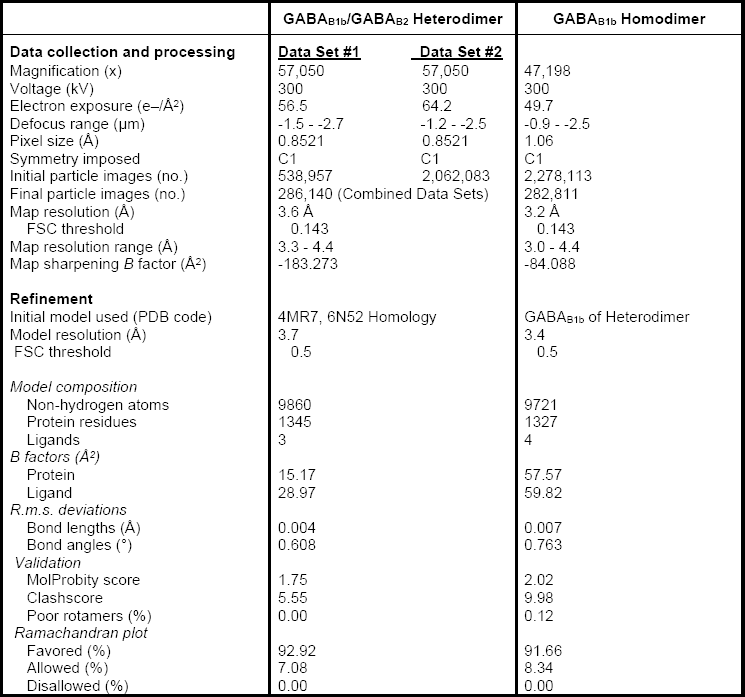
Cryo-EM data collection, refinement and validation statistics.

**Supplemental Table 2.**
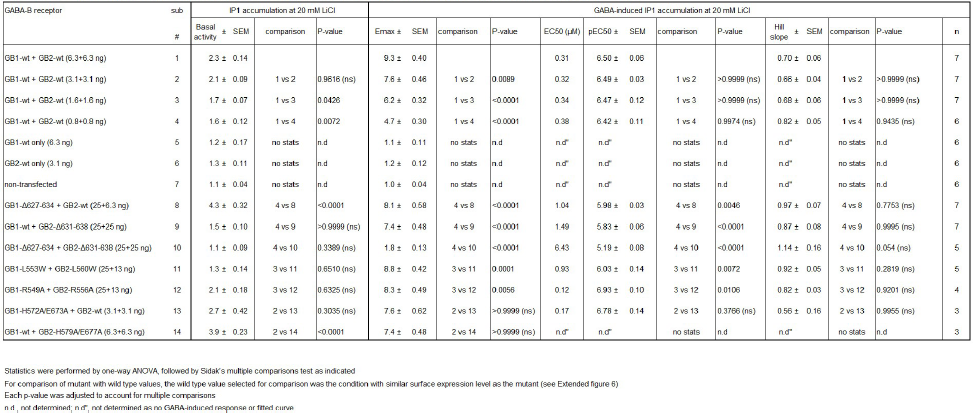
Basal, Emax and GABA potency extimates of GABA-B ECL2 deletion mutants in HEK293 cells co-transfected with Gqo5.

